# Phase-based coordination of hippocampal and neocortical oscillations during human sleep

**DOI:** 10.1101/745745

**Authors:** Roy Cox, Theodor Rüber, Bernhard P Staresina, Juergen Fell

**Affiliations:** Department of Epileptology, University of Bonn, 53127 Bonn, Germany; Epilepsy Center Frankfurt Rhine-Main, Department of Neurology, Goethe University Frankfurt, 60590 Frankfurt am Main, Germany; Center for Personalized Translational Epilepsy Research (CePTER), Goethe University Frankfurt, 60590 Frankfurt am Main, Germany; School of Psychology, University of Birmingham, B15 2TT Birmingham, United Kingdom

## Abstract

During sleep, new memories undergo a gradual transfer from the hippocampus (HPC) to the neocortex (NC). Precisely timed neural oscillations interacting within and between these brain structures are thought to mediate this sleep-dependent memory consolidation, but exactly which sleep oscillations instantiate the HPC-NC dialog, and via what mechanisms, remains elusive. Employing invasive electroencephalography in ten neurosurgical patients across a full night of sleep, we identified three broad classes of phase-based HPC-NC communication. First, we observed interregional phase synchrony for non-rapid eye movement (NREM) spindles, N2 and rapid eye movement (REM) theta, and N3 beta activity. Second, and most intriguingly, we found asymmetrical N3 cross-frequency phase-amplitude coupling between HPC SOs and NC activity spanning the delta to high-gamma/ripple bands, but not in the opposite direction. Lastly, N2 theta and NREM spindle synchrony were themselves modulated by HPC SOs. These novel forms of phase-based interregional communication emphasize the role of HPC SOs in the HPC-NC dialog, and may offer a physiological basis for the sleep-dependent reorganization of mnemonic content.

## Introduction

A long-standing question in cognitive neuroscience asks how initially fragile episodic memories are transformed into lasting representations. Theoretical accounts postulate that this process involves a protracted transfer of memories from the hippocampus (HPC) to neocortical (NC) domains (1–3), with a large body of lesion (4,5), and neuroimaging (6,7) findings supporting this notion. One NC area of particular interest is the lateral temporal cortex, a convergence zone involved in long-term memory storage (8,9), representing higher order visual, categorical, and semantic concepts (10–12).

Intriguingly, sleep leads to more stable and better integrated episodic memories (13,14), suggesting a pivotal role for this brain state in the systems-level reorganization of memory traces (15). Neural oscillations, especially non-rapid eye movement (NREM) neocortical slow oscillations (SOs; 0.5–1 Hz), thalamocortical sleep spindles (12.5–16 Hz), and hippocampal ripples (80–100 Hz), are widely held to mediate the HPC-NC memory transfer and consolidation process (16–21), particularly given the presence of both SOs and spindles in HPC (22–26). Moreover, various other spectral components exist in electrophysiological recordings of human sleep, with recent evidence suggesting potential roles for theta (4–8 Hz) in NREM (27,28) and rapid eye movement (REM) (29,30) memory processing, complicating the question of which oscillatory rhythms instantiate the HPC-NC dialog.

Oscillatory phase (i.e., the relative position along the oscillatory cycle) has a critical influence on neuronal excitability and activity (31), thereby offering a precise temporal scaffold for orchestrating neural processing within and across brain structures (32,33). As such, oscillatory phase coordination between HPC and NC is a prime candidate mechanism for sleep-dependent information exchange between these areas. However, various forms of phase coupling may be distinguished, and phase-based HPC-NC interactions during sleep could be implemented in at least three (non-mutually exclusive) ways.

First, consistent oscillatory phase locking between brain regions at the same frequency is thought to enable effective communication between the underlying neuronal groups (34). Phase synchrony during sleep has been reported between neocortical regions for various frequency bands (35), including the spindle (36,37) and gamma (38) ranges, and between HPC and prefrontal areas in the spindle range (39). Whether similar phenomena exist between human HPC and non-frontal NC areas, for which frequency bands, and in which sleep stages, has not been examined.

Beside potential phase coupling *within* frequency bands, NREM sleep oscillations are also temporally organized *across* frequency bands. Such cross-frequency phase-amplitude coupling (PAC) is thought to enable brain communication across multiple spatiotemporal scales (40,41). Local PAC among SOs, spindles, and ripples has been well characterized for various brain structures including HPC (24,26,36,42–47), and is considered a fundamental building block of memory consolidation theories (48). However, local PAC exists for other frequency pairs (26), with SOs exerting particularly powerful drives not only over spindle and ripple activity, but also over delta (49), theta (50), and gamma (36,38) components. Extending the notion of local PAC to cross-regional interactions, the phase of a slower rhythm in one brain structure may modulate expression of faster activity at the other site (24,47,51), thus constituting a second potential form of HPC-NC communication.

Third, interregional phase synchronization within a frequency band might itself be modulated by the phase of a slower rhythm, as shown for the SO-phase-dependent coordination of spindle synchrony in neocortical networks (36). Whether analogous SO-based modulation of phase synchronization exists between HPC and NC, and if so, for which frequency components, remains unexplored.

Here, we examined intracranial electrophysiological activity in a sample of 10 presurgical epilepsy patients during light NREM (N2), deep NREM (N3) and REM sleep. Specifically, we focused on HPC and lateral temporal cortex as a site relevant for long-term memory storage. We hypothesized that these areas exhibit interregional phase coordination, which could manifest in any or all of the three aforementioned forms of coupling. Given both the theoretical importance of nested SO-spindle-ripple activity (52), and inconclusive evidence regarding the directionality of HPC-NC coupling (22,25,39,53–57), we were particularly interested in whether HPC and NC SOs or spindles modulate faster activity at the other brain site, and if so, whether these effects are direction-dependent. Moreover, we considered a wide 0.5–200 Hz frequency range to allow potential identification of oscillatory communication lines outside the canonical SO-spindle-ripple framework. Using this approach, we identified several novel forms of phase-based HPC-NC communication centered on SO, spindle, theta, and beta activity, thereby offering a potential neurobiological substrate for sleep-dependent memory consolidation.

## Results

We analyzed overnight invasive electroencephalography (EEG) from the hippocampus (HPC) and lateral temporal neocortex (NC) in a sample of 10 epilepsy patients during N2, N3 and REM sleep. Polysomnography-based sleep architecture was in line with healthy sleep (Table S1). Only intracranial contacts from the non-pathological hemisphere were used, as evidenced by clinical monitoring. Electrode locations are shown in Fig. 1 and Table S2.

**Figure 1.**
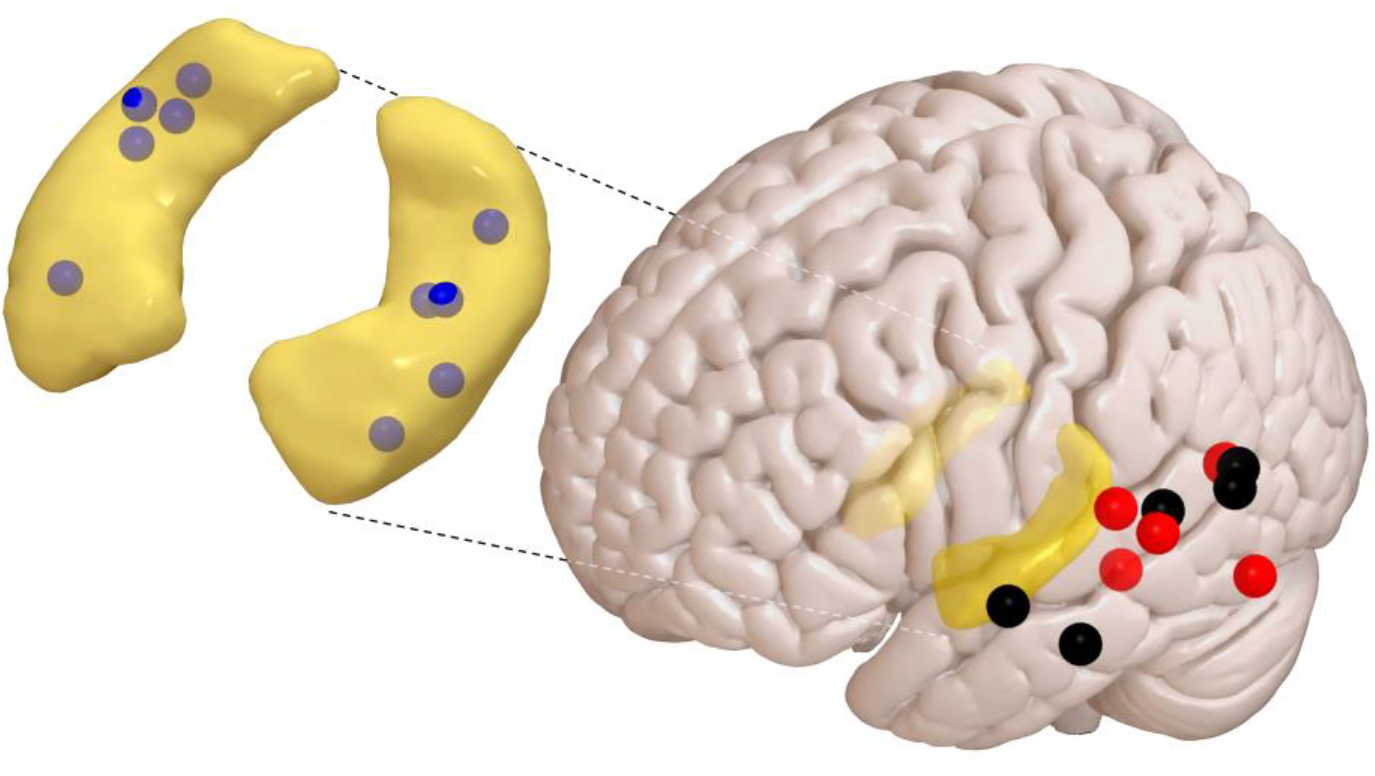
Group-level distribution of electrodes in hippocampus and on neocortical surface. Outline of hippocampi in yellow, hippocampal depth electrodes in blue, subdural neocortical electrodes in red (left hemisphere) and black (flipped from right hemisphere). Electrode contacts not to scale.

### Spindle, theta, and beta phase synchronization between hippocampus and neocortex

Following inspection of raw traces with spectrograms (Fig. S1), and power spectra (Fig. S2), we evaluated whether, and to what degree, oscillatory signals in HPC and NC show phase coordination within frequency bands. Using the weighted phase lag index (wPLI: a metric minimally sensitive to common neural sources (58)), we observed that raw wPLI showed a general decrease with frequency (Fig. 2A), with slower rhythms showing stronger synchrony than fast oscillations, as typically observed (59). However, clear departures from this downward trend appeared for three main frequency bands. First, phase synchrony was enhanced in the spindle range (13.6-15.3 Hz) during N2 and especially N3 sleep, consistent with findings from other brain sites (36,39). Surprisingly, however, synchronization enhancements were also observed in the theta range (7.4-8.3 Hz) during N2 and REM, and in the beta range (28 Hz) during N3.

To determine whether phase coupling in these or any other frequency bands was beyond chance levels, we z-scored raw wPLI values with respect to time-shifted surrogate distributions. This procedure essentially removed the downward trend, while retaining the aforementioned theta, spindle, and beta peaks (Fig. 2B). Comparing wPLI_Z_ values to zero (one-tailed Wilcoxon signed rank test with False Discovery Rate [FDR] correction; significant ranges indicated by colored bars at bottom of Fig. 2B) yielded above-chance HPC-NC phase coordination in the theta range for N2 (P_uncorrected_<0.007) and REM (P_uncorrected_<0.03). Reliable spindle connectivity was seen for N3 (P_uncorrected_<0.002) and REM sleep (P_uncorrected_ <0.001). Phase synchrony for the N2 spindle (P_uncorrected_=0.12) and N3 beta bands (P_uncorrected_=0.01) were not significant after correction, but both effects were clearly seen for individual patients (Fig. S3ABC). Other group-level effects of above-zero coupling were observed in the SO/delta range during REM (P_uncorrected_<0.05), and for N2 delta (P_uncorrected_<0.003). Interestingly, no SO-based connectivity was seen during N2 or N3, suggesting that SO phase relations between HPC and NC are relatively variable (25,39).

**Figure 2.**
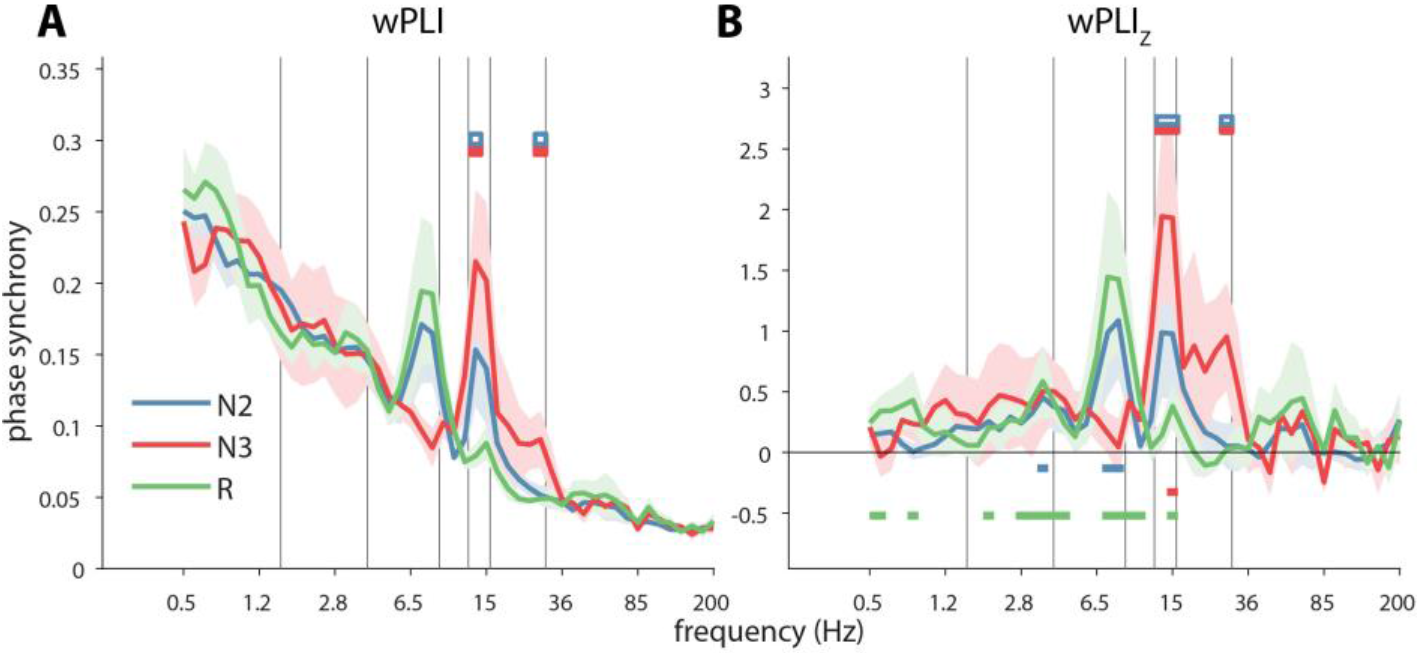
Phase synchrony between hippocampus and necortex. Group-level connectivity profiles for (unnormalized) wPLI (A) and (normalized) wPLIZ (B). Horizontal color bars at top of both panels indicate significant pairwise stage differences (two-tailed Wilcoxon signed rank tests with FDR correction). Filled color reflects stage with greater connectivity. Horizontal color bars at bottom of panel (B) indicate above-chance connectivity (one-tailed Wilcoxon signed rank tests versus zero with FDR correction). Error shading: standard error of the mean across patients. Gray vertical lines at 1.5, 4, 9, 12.5, 16 and 30 Hz indicate approximate boundaries between SO, delta, theta, slow spindle, fast spindle, beta, and faster activity.

Sleep stage comparisons for both raw and normalized phase synchrony indicated reliable N2/N3 differences in the spindle and beta frequency bands (colored bars at top of Fig. 2, two-tailed Wilcoxon signed rank test with FDR correction, all P_uncorrected_<0.006), providing further support for the existence of HPC-NC communication in these bands during N3. Finally, various control analyses showed that phase synchrony was not systematically related to power (SI Results). Overall, these findings indicate a precise phase-based coordination between HPC and NC rhythms, primarily in the spindle, theta, and beta ranges, thus signifying the existence of communication lines beyond the canonical NREM oscillators.

### Cross-frequency coupling of neocortical activity to hippocampal slow oscillations

Next, we turned our attention to interactions between, rather than within, frequency bands. We quantified cross-frequency coupling using the debiased phase-amplitude coupling metric (dPAC: a metric correcting for potential non-sinusoidality of the phase-providing frequency (60)). These values were further z-scored with respect to surrogate distributions. The resulting metric (dPAC_Z_) signifies the degree to which activity at a faster frequency is non-uniformly distributed across the phase of a slower frequency.

Following analyses of local PAC within HPC and NC separately (SI Results and Fig. S4-S6), we asked whether the oscillatory phase in one brain area could modulate faster activity in the other region. Assessing whether the phase of HPC rhythms coordinates faster activity in NC (“HPC-NC PAC”), we found that HPC SOs (0.5–1 Hz) robustly orchestrate the expression of faster activity in NC during N3 sleep (Fig. 3A). Specifically, distinct hotspots were found for modulated frequencies in the delta (maximum: 3.5 Hz), theta (6.5 Hz), beta/low-gamma (32 Hz), and high-gamma/ripple (85 Hz) ranges (white arrows in Fig. 3A). No systematic cross-regional modulation of neocortical activity by the hippocampal phase was observed during N2 or REM sleep. Interestingly, and in stark contrast to the robust modulation of NC activity by HPC SOs, the NC SO phase did not reliably coordinate faster HPC dynamics for any frequency band or sleep stage, nor did the phase of any other NC frequency modulate HPC activity (“NC-HPC PAC”, Fig. 3B).

**Figure 3.**
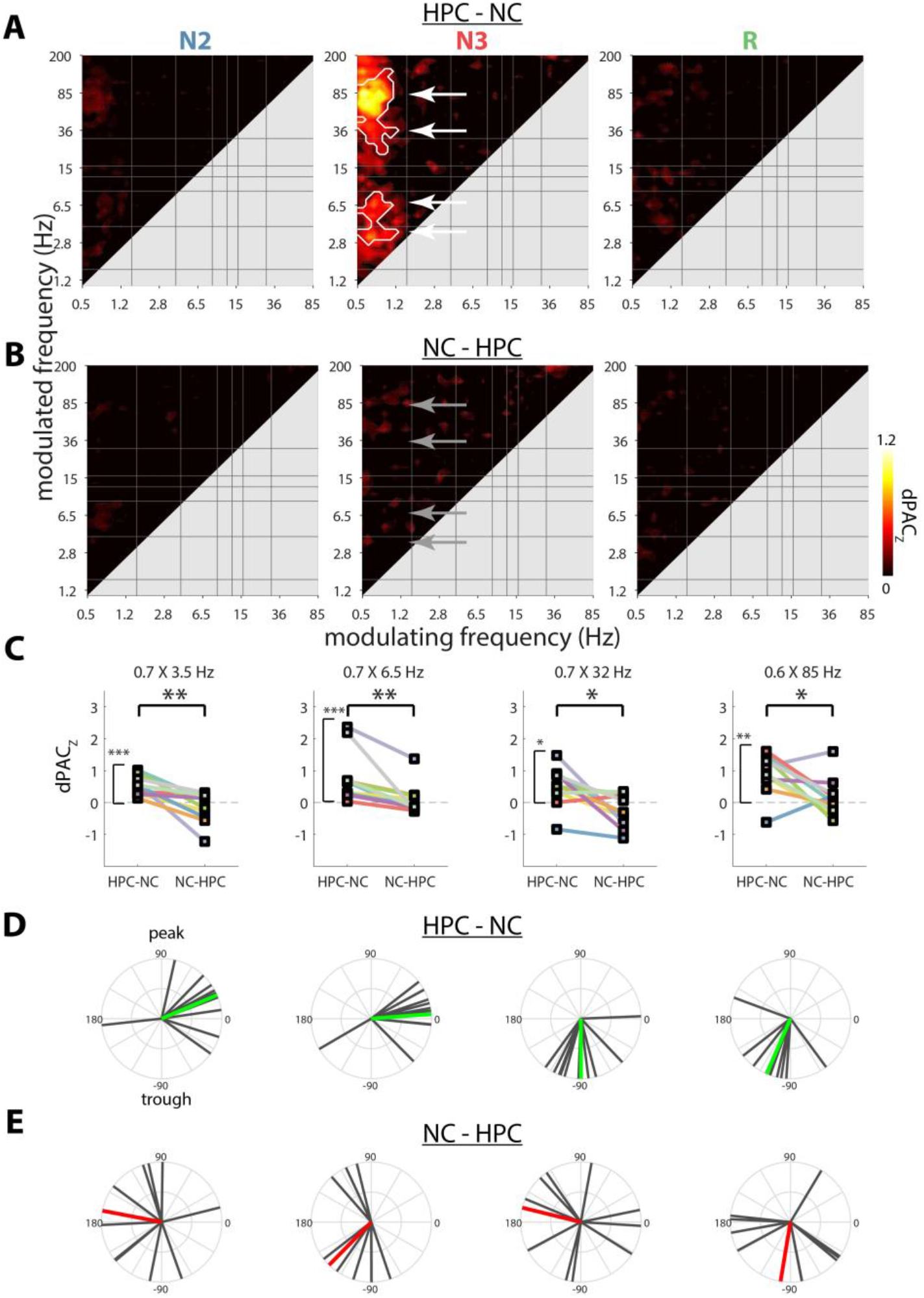
Cross-frequency coupling between hippocampus and neocortex. Coupling strengths for HPC-NC (A) and NC-HPC PAC (B). White outlines indicate clusters of significantly greater than zero coupling across patients (cluster-based permutation test). (C) Comparisons of HPC-NC and NC-HPC PAC for each SO-based N3 cluster (indicated in panels A and B with arrows). * P<0.05, ** P<0.01, *** P<0.001 (Wilcoxon signed rank test, uncorrected). (D and E) SO phase (with respect to sine wave) at which faster activity is maximally expressed across patients for HPC-NC (D) and NC-HPC (E) PAC. Colored lines indicate group averages, with green indicating significant (P<0.05) deviations from uniformity, and red nonsignificance.

We further investigated this apparent asymmetry in how the N3 SO rhythm coordinates distant activity in various manners. First, we extracted individuals’ dPAC_Z_ values for the four frequency pairs showing maximum HPC-NC group effects (white arrows in Fig. 3A), along with their opposite direction counterparts (gray arrows in Fig. 3B). Unsurprisingly, coupling strength for each selected frequency pair was significantly greater than zero for the HPC-NC direction (one-tailed Wilcoxon signed rank test with FDR, all P_corrected_<0.02). In contrast, no above-chance coupling was observed in the opposite NC-HPC direction (all P_uncorrected_>0.31). Moreover, directional comparisons indicated that interregional PAC was systematically greater for HPC-NC vs. NC-HPC coupling for each frequency pair (two-tailed Wilcoxon signed rank test: all P_corrected_<0.03), as further illustrated in Fig. 3C. We also directly compared the full interregional coupling profiles of Fig. 3A and B, yielding a highly similar pattern of enhanced HPC-NC vs. NC-HPC coupling centered on the N3 SO-band (Fig. S7). Of note, individual profiles of interregional coupling were highly consistent with these group-level findings (Fig. S8).

Second, we considered, for the SO-based frequency pairs of N3, the precise phase at which distant fast activity was maximally expressed. For HPC-NC PAC (Fig. 3D), phase distributions deviated substantially from uniformity for each of the four clusters (Rayleigh test for uniformity with FDR, all P_corrected_<0.008), indicating that fast NC activity is preferentially expressed at similar phases of the HPC SO across patients. Specifically, delta and theta activity occurred around the negative-to-positive zero-crossing, while the low-gamma and high-gamma/ripple bands showed maximal activity in the SO trough (likely reflecting the physiological up state (22,26)). In contrast, HPC fast activity was not consistently expressed in a particular phase range of the NC SO (all P_uncorrected_>0.09, Fig. 3E), consistent with the lack of coupling reported in the previous paragraph.

Finally, we directly compared interregional HPC-NC PAC (as shown in Fig. 3A) to local PAC within each brain structure (as shown in Fig. S4AB). For both HPC and NC, cross-frequency interactions were generally stronger within than between brain structures, as shown in Fig. 4. Specifically, the phase of HPC delta (1.5–4 Hz) organized spindle/beta and ripple activity more strongly within local HPC than in distant NC, most prominently during N3 (Fig. 4A). This delta-ripple effect is consistent with sharp-wave-ripple complexes (61). Interestingly, the N3 modulation of faster activity by the HPC SO was spared from these effects of enhanced local vs. interregional PAC (red oval in Fig. 4A), suggesting that HPC SOs are equally capable of modulating faster components in local and distant brain sites. In contrast, fast NC activity in the spindle-to-high-gamma bands during N3 was coordinated more robustly by the local NC SO than the HPC SO (black arrow in Fig. 4B). These observations could indicate that while fast NC activity is under the control of HPC SOs, local SOs still exert a stronger influence. While some local vs. interregional differences were also seen for N2 and REM, these effects should be interpreted cautiously, since no systematic interregional PAC was seen for these sleep stages (Fig. 3A).

**Figure 4.**
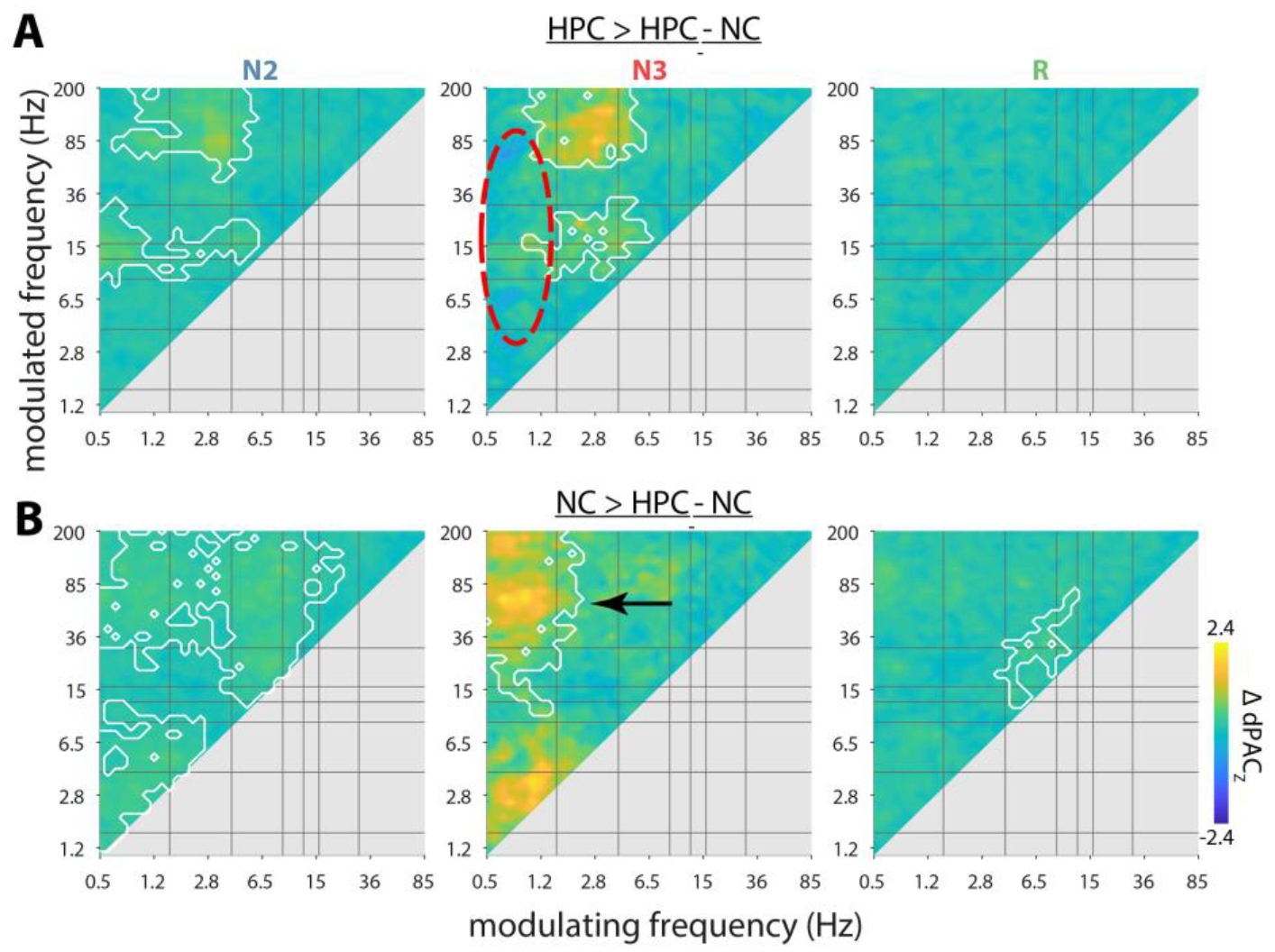
Differences between local and interregional cross-frequency coupling. Coupling strength differences for HPC versus HPC-NC (A) and NC versus HPC-NC (B). White outlines indicate clusters of significantly greater same-site than cross-site coupling (cluster-based permutation). No clusters with greater cross-site than same-site coupling were observed.

Overall, these findings indicate that the HPC SO phase is capable of coordinating the expression of faster activity in NC regions during N3 sleep, whereas the reverse NC-HPC modulation does not occur. Given these observations, as well as the within-frequency synchronization for faster rhythms (Fig. 2), an intriguing possibility is that SO rhythms also affect interregional phase synchronization. We address this question next.

### Modulation of interregional phase synchronization by hippocampal slow oscillations

As a final potential form of phase-based HPC-NC communication, we asked whether within-frequency phase synchronization for faster frequencies could vary as a function of a slower oscillatory phase. We computed HPC-NC wPLI for each modulated frequency as a function of the phase (18 bins) of each slower frequency in either HPC or NC. We then determined a modulation index (MI) (62) for each frequency pair, indicating the degree to which wPLI values are non-uniformly distributed across the cycle of a slower frequency, and further normalized MI with respect to surrogate distributions. (Due to methodological considerations related to data length, two patients were excluded from N3 analyses.)

Intriguingly, these analyses revealed a strong organizing influence from the HPC SO on interregional phase synchronization (Fig. 5A). Specifically, HPC-NC theta synchrony was reliably modulated by HPC SOs during N2, whereas spindle synchrony was coordinated by both N2 and N3 HPC SOs, similar to scalp findings (36). Note that these theta and spindle effects overlap well with the frequency bands showing interregional synchrony in Fig. 2. No other frequency bands showed synchronization modulations in relation to the phase of any other HPC oscillation, for neither NREM nor REM sleep. Moreover, we observed no statistically reliable modulation of phase synchronization by the NC phase, neither for SOs nor other frequency bands (Fig. 5B), although we note that hotspots suggestive of SO-spindle effects were visually apparent.

**Figure 5.**
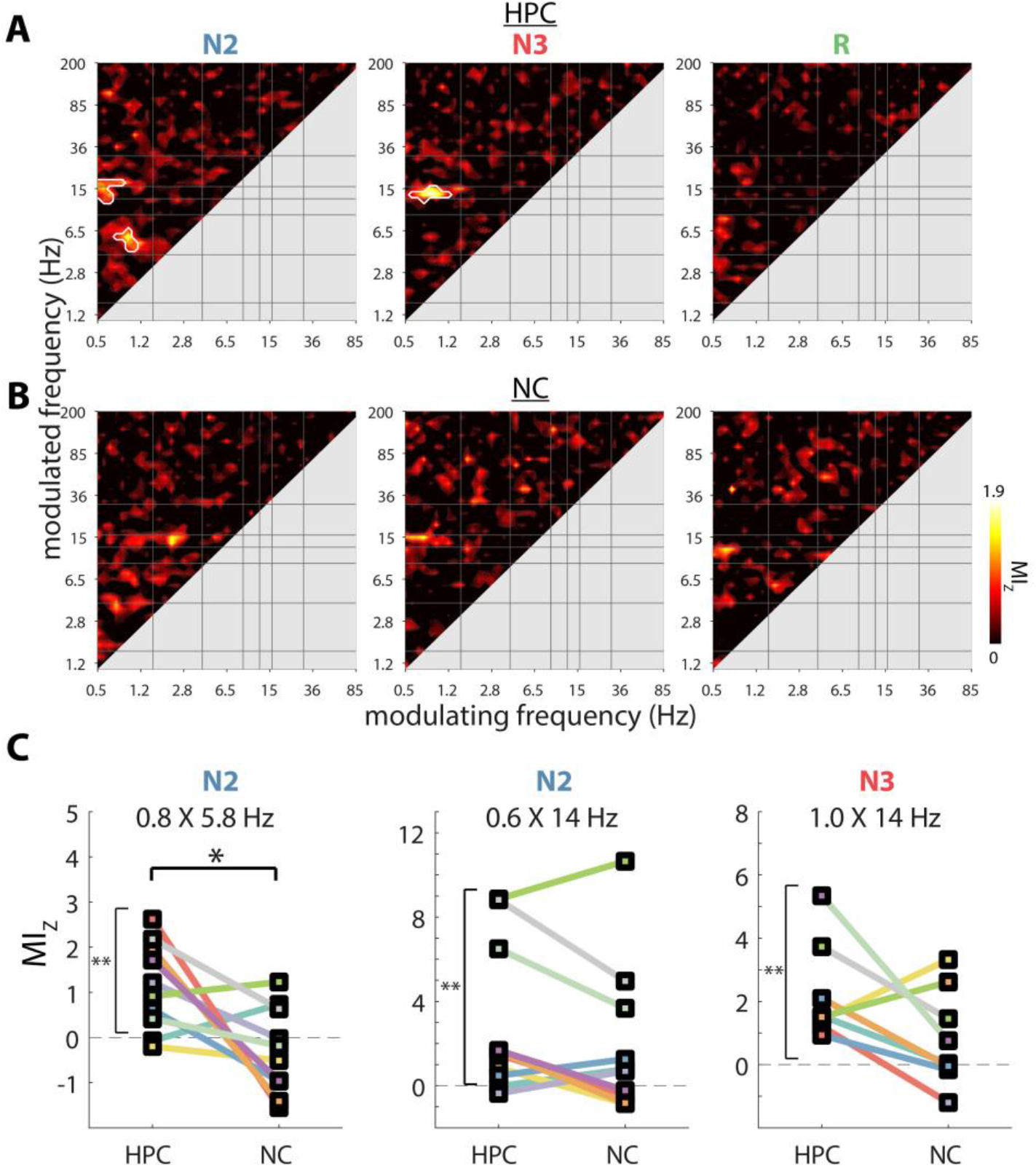
Cross-frequency modulation of phase synchronization between hippocampus and neocortex. Normalized modulation indices (MIZ) for phase of HPC (A) and NC (B). White outlines indicate clusters of significantly greater than zero modulation across patients (cluster-based permutation test). (C) Comparisons of HPC− and NC-phase-dependent modulation of phase synchronization for each SO-based cluster. * P<0.05, ** P<0.01, *** P<0.001 (Wilcoxon signed rank test, uncorrected). N=8 for N3, N=10 for N2 and REM.

Similar to our approach for cross-regional PAC, we extracted patients MI_Z_ values for each of the three significant clusters of Fig. 5A. As visualized in Fig. 5C, these were all (trivially) significantly greater than zero (one-tailed Wilcoxon signed rank test with FDR, all P_corrected_<0.005), whereas their counterparts in the other brain region did not differ reliably from zero (all P_uncorrected_>0.09). Direct comparisons between HPC− and NC-based modulation of phase synchronization for these frequency pairs revealed a reliably greater HPC vs. NC influence of N2 SOs on theta synchrony (two-tailed Wilcoxon signed rank test with FDR, P_corrected_=0.08, P_uncorrected_=0.03). In contrast, while spindle synchrony also appeared to be more robustly modulated by the phase of HPC rather than NC SOs (Fig. 5C), these effects did not reach significance (both P_uncorrected_=0.11). Finally, we directly compared the full HPC and NC profiles of Fig. 5AB, but no significant clusters emerged (Fig. S9).

In sum, the phase of SO activity orchestrates interregional phase synchronization in both the theta and spindle frequency bands, with HPC SOs having a particularly strong impact on theta synchrony. These findings establish another major form of phase-based HPC-NC coordination, potentially contributing to systems-level memory reorganization.

## Discussion

Communication between the hippocampus and neocortex during sleep is considered a cornerstone of theories of memory consolidation, but exactly how these interactions are instantiated in the human brain has remained unclear. In line with the notion that oscillatory phase is critically involved in binding distant but functionally related neural populations (32), we observed systematic i) within-frequency phase synchronization, ii) cross-frequency phase-amplitude coupling, and iii) cross-frequency modulation of within-frequency phase synchronization, thereby uncovering several previously unknown modes of interregional HPC-NC communication. A particularly prominent role emerged for HPC SOs, coordinating both the expression of, and synchronization with, faster NC activity.

### Phase synchrony between hippocampus and neocortex

As a first major form of phase-based interregional communication, we observed within-frequency theta, spindle, and beta phase synchronization between HPC and NC (Fig. 2), thus reflecting precise oscillatory coordination on a cycle-by-cycle basis for these frequency bands. Consistent phase relations between brain areas affect the relative timing of neuronal spikes, thereby enabling communication and plasticity (34,63).

Sleep spindles are closely tied to memory and plasticity (17,18,64,65), and show widespread phase synchronization in neocortical networks (36). Here, we extend these observations of NREM spindle synchrony to include dynamics between HPC and lateral temporal cortex, similar to recent observations between HPC and prefrontal areas (39). Thus, the precise coordination of spindle activity across HPC and distributed neocortical areas may offer a potential mechanism to transiently reactivate distributed memory traces, and thereby contribute to NREM-dependent memory consolidation.

Surprisingly, similar observations of HPC-NC phase synchrony were made for N2 and REM theta. These findings may offer a physiological basis for recent work demonstrating a role for NREM theta in memory consolidation (27,28). In contrast, interregional HPC-NC theta synchrony during REM sleep could form a neurobiological basis for associations between REM theta and the regulation and consolidation of emotional content (29,30). Combined with similar findings of REM theta connectivity between prefrontal and cingulate areas (66), theta rhythms thus appear to be coordinated across widespread brain areas. Of note, the observed REM theta synchrony contrasts with a study reporting no REM theta coherence between HPC and NC (67). However, that observation was based on only two patients, providing limited opportunities to detect effects that may not be present in all individuals, as we also observed (e.g., Fig. S3A).

Another unexpected spectral component to exhibit phase synchrony was observed in the beta band during N3. While not present in every individual (and therefore weaker at the group level), a beta effect clearly distinct from the spindle frequency band could be discerned on an individual basis (Fig. S3CF). Thus, beta activity may reflect another spectral band of interest regarding sleep-based communication.

Of note, NREM synchrony in each of the aforementioned frequency bands varied with sleep depth. While the reason for differential connectivity during N2 and N3 is unclear, these findings underscore the need to consider these sleep stages separately. The absence of reliable phase synchrony in other frequency bands also deserves mention. Although raw wPLI was greatest for slower rhythms (Fig. 2A), SO synchronization after surrogate-based normalization was absent during N2 and N3 (Fig. 2B), suggesting that HPC and NC SOs show variable phase relations (25,39). Similarly, ripple band activity was not reliably synchronized between brain structures, consistent with analogous findings of low co-occurrence of neocortical gamma events (38). However, we note that phase synchrony profiles differed between individuals, with synchrony in SO, ripple, and other frequency bands sometimes reliably expressed on an individual basis (Fig. S3ABC).

### Cross-frequency phase-amplitude coupling between hippocampus and neocortex

As a second major form of oscillatory HPC-NC coordination, we observed systematic interregional cross-frequency phase-amplitude coupling. These effects were restricted to a governing role of HPC SOs over NC activity spanning the delta, theta, low gamma, and ripple ranges (Fig. 3A). The opposite pattern, whereby the phase of NC oscillations coordinates HPC activity, for either SOs or other frequencies, was not seen (Fig. 3BC). Similarly, the preferred SO phase at which faster activity was expressed was highly consistent across patients for HPC-NC, but not NC-HPC PAC (Fig. 3DE). Of note, this asymmetry is consistent with the notion of independent HPC and NC SO dynamics, as suggested by the lack of NREM SO phase synchrony.

Although our metric of interregional PAC does not contain directional information *per se*, it is widely assumed that it is the phase of the slower frequency that modulates faster activity, rather than the other way around (40,41). While SOs and their coordination of faster activity are typically viewed as NC phenomena (Fig. S4B), similar dynamics within HPC are now well established (Fig. S4A) (22,24–26,53,68,69). As such, our findings suggest a driving force of HPC SOs on NC activity, co-determining NC activity in various faster frequency bands. These effects may stem from surges of local activity associated with HPC up states being transmitted to post-synaptic targets and eventually reaching NC. Indeed, while faster activity was typically modulated more strongly by local than distant slower rhythms, HPC SO activity coordinated local and NC faster activity to similar extents (Fig. 4A), potentially fostering more efficient HPC-NC information exchange.

We did not observe systematic cross-regional HPC-NC PAC for modulating rhythms beyond SOs, although we did find such examples on an individual basis (e.g., HPC theta modulating NC beta/gamma/high gamma activity, Fig. S8A, p7, N2). This general lack of HPC-NC PAC beyond SOs is noteworthy given that many additional frequency pairs were coupled locally in HPC and NC (Fig. S4). These findings indicate that cross-regional and local PAC are at least partially dissociated, which is further supported by the observed asymmetry between HPC-NC and NC-HPC PAC. Importantly, these findings also alleviate concerns that cross-regional PAC is due to volume conduction, whereby modulating, modulated, or both signal components primarily reflect activity from the other brain site.

The lack of systematic NC-HPC PAC in our data may appear at odds with previous observations of HPC spindles and ripples coupled to NC SOs and spindles, respectively (24,47,51). However, previous studies observing NC-HPC PAC assessed NC activity with non-invasive scalp electrodes that aggregate activity over large spatial domains, thus reflecting common signals with relatively powerful drives. In contrast, the localized NC activity we considered here constitutes only a tiny fraction of all NC activity and may therefore exert a more limited influence on HPC activity (also see relative dissociation of scalp and NC signals in Fig. S1).

### Cross-frequency modulation of phase synchronization

The third and final form of oscillatory HPC-NC interaction we observed was the modulation of within-frequency phase synchronization by the phase of slower rhythms. Most prominently, the HPC SO phase had a robust influence on the degree of N2 theta and NREM spindle synchrony (Fig. 5A), matching the sleep stages where these forms of synchrony were apparent (Fig. 2).

The gating of spindle synchrony by HPC SOs is highly consistent with similar observations of SO-modulated spindle synchrony in scalp data (36), and compatible with findings of enhanced HPC-prefrontal spindle synchrony for spindles coupled vs. uncoupled to frontal SOs (39). While we did not see unambiguous evidence that NC SOs impose a similar modulation on spindle synchrony, modulation strengths also did not differ reliably between HPC and NC. Hence, strong conclusions regarding whether HPC or NC SOs most effectively affect spindle synchrony are presently not warranted.

In stark contrast, N2 theta synchrony depended to a greater extent on HPC than NC SOs. Intriguingly, this effect appears to be separate from the enhanced HPC-NC vs. NC-HPC modulation of theta amplitude by SOs, which occurred in N3 rather than N2. While the reason for this dissociation is unclear, both effects are in agreement that interregional theta dynamics are modulated most effectively by HPC rather than NC SOs.

### Relevance and limitations

A major unresolved issue concerns the directionality of HPC-NC dynamics during sleep. Prior empirical evidence has been mixed, pointing towards HPC-NC (56,70), NC-HPC (22,54,71), or more elaborate bidirectional paths (39,55,57). Both our findings of HPC SOs coordinating NC faster activity but not *vice versa*, and of stronger modulations of theta phase synchrony by HPC SOs than NC SOs, are most consistent with the notion of HPC to NC directionality, as suggested by classical theoretical (15) and computational (72) models. Based on these results, we propose that HPC SOs may influence plasticity not only within HPC but also in NC circuits (53).

We examined communication between HPC and lateral temporal cortex, a neocortical region that has not received much attention in the context of sleep-dependent memory consolidation. Although precise electrode placement varied across patients (Fig. 1, Table S2), the role of lateral temporal cortex in long-term memory storage (8,9) makes it well-suited for studying HPC-NC interactions. Future work should address whether the reported forms of phase coordination apply more broadly to other NC sites.

Although generalizing from epileptic to healthy populations poses a risk, sleep architecture was in line with healthy sleep (Table S1). Moreover, we employed a rigorous artifact rejection protocol, and only considered electrodes on the non-pathological side, making it unlikely our results are due to epileptogenic activity. In the present approach, measures of oscillatory coordination were calculated over continuous data. This contrasts with discrete approaches where analyses are contingent on the presence of specific waveforms. Given that our approach identified various expected phenomena of local PAC (e.g., SO-spindle, spindle-ripple; Fig. S4 and S6), we do not believe this methodological choice poses a major concern. That said, future work may scrutinize individual waveforms to fully understand the origin of each of the observed effects (73).

### Conclusion

The present observations establish an important prerequisite for memory consolidation theories postulating a sleep-dependent HPC-NC dialog (2,13,14). More specifically, the identified forms of phase coordination draw attention not only to SOs and spindles, but also to theta and beta activity. Most intriguingly, the asymmetrical coordination of NC activity and HPC-NC phase synchronization by HPC SOs suggests that HPC may play a larger orchestrating role in information exchange during sleep than previously thought. Overall, these findings refine our knowledge of human HPC-NC interactions and offer novel opportunities to understand the determinants of sleep-dependent memory consolidation in health and disease.

## Materials and Methods

### Participants

We analyzed archival electrophysiological sleep data in a sample of 10 (6 male) patients suffering from pharmaco-resistant epilepsy (age: 36.6 ± 14.8 yrs, range: 22–62). This sample overlaps with ones reported previously (24,26,54). Local aspects of the HPC data are described in detail elsewhere (26), but are summarily included here both because of different patient and electrode inclusion criteria and to provide a comprehensive perspective on HPC-NC oscillations. Patients had been epileptic for 22.5 ± 11.0 yrs (range: 10–49) and were receiving anticonvulsive medication at the moment of recording. All patients gave informed consent, the study was conducted according to the Declaration of Helsinki, and was approved by the ethics committee of the Medical Faculty of the University of Bonn.

### Data acquisition

Electrophysiological monitoring was performed with a combination of depth and subdural strip/grid electrodes. HPC depth electrodes (AD-Tech, Racine, WI, USA) containing 8–10 cylindrical platinum-iridium contacts (length: 1.6 mm; diameter: 1.3 mm; center-to-center inter-contact distance: 4.5 mm) were stereotactically implanted. Implantations were done either bilaterally (n=7) or unilaterally (n=3), and either along the longitudinal HPC axis via the occipital lobe (n=9) or along a medial-lateral axis via temporal cortex (n=1). Stainless steel subdural strip/grid electrodes were of variable size with contact diameters of 4 mm and center-to-center spacing of 10 mm, and placed over various neocortical areas according to clinical criteria. Anatomical labels of each electrode were determined based on pre− and post-implantation magnetic resonance image (MRI) scans by an experienced physician (TR), as described previously (26).

A single grey matter HPC electrode and a single NC electrode from lateral temporal cortex were selected for each patient. As reported previously for HPC, and here additionally seen for NC, within-patient spectral profiles varied between adjacent contacts. Following previous approaches (24) and the hypothesized central role of spindles in HPC-NC communication, the contact with highest NREM spindle power was chosen at both brain sites. MNI electrode locations for each patient are indicated in Table S2, and were visualized using Surf Ice (https://www.nitrc.org/projects/surfice/) to generate Fig. 1. For HPC, fast spindle peaks were visible for all patients. For NC, 7 of 10 patients showed fast spindle peaks, one showed a slow spindle peak, and two did not exhibit noticeable spindle peaks. Additional non-invasive signals were recorded from the scalp (Cz, C3, C4, Oz, A1, A2; plus T5 and T6 in 8 patients), the outer canthi of the eyes for electrooculography (EOG), and chin for electromyography (EMG). All signals were sampled at 1 kHz (Stellate GmbH, Munich, Germany) with hardware low− and high-pass filters at 0.01 and 300 Hz respectively, using an average-mastoid reference. Offline sleep scoring was done in 20 s epochs based on scalp EEG, EOG, and EMG signals in accordance with Rechtschaffen and Kales criteria (74). Stages S3 and S4 were combined into a single N3 stage following the more recent criteria of the American Academy of Sleep Medicine (75).

### Preprocessing and artifact rejection

All data processing and analysis was performed in Matlab (the Mathworks, Natick, MA), using custom routines and EEGLAB functionality (76). Preprocessing and artifact rejection details are identical to our previous report (26). Briefly, data were high-pass (0.3 Hz) and notch (50 Hz and harmonics up to 300 Hz) filtered, and channel-specific thresholds (z-score > 6) of signal gradient and high-frequency (>250 Hz) activity were applied to detect and exclude epileptogenic activity. Artifact-free data “trials” of at least 3 s were kept for subsequent processing, resulting in a total of 78.1 ± 30.8 (N2), 21.7 ± 17.8 (N3), and 44.5 ± 23.7 min (REM) of usable data.

### Time-frequency decomposition

Data were decomposed with a family of complex Morlet wavelets. Each trial was extended with 5 s on either side to minimize edge artifacts. Wavelets were defined in terms of desired temporal resolution according to:

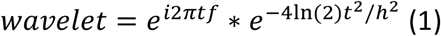

where *i* is the imaginary operator, *t* is time in seconds, *f* is frequency (50 logarithmically spaced frequencies between 0.5 and 200 Hz), *ln* is the natural logarithm, and *h* is temporal resolution (full-width at half-maximum; FWHM) in seconds (77). We set *h* to be logarithmically spaced between 3 s (at 0.5 Hz) and 0.025 s (at 200 Hz), resulting in FWHM spectral resolutions of 0.3 and 35 Hz, respectively. Trial padding was trimmed from the convolution result, which was subsequently downsampled by a factor four to reduce the amount of data. We normalized phase-based metrics using surrogate approaches (see below). To make surrogate distributions independent of variable numbers and durations of trials, we first concatenated the convolution result of all trials of a given sleep stage, and then segmented them into 60 s fragments (discarding the final, incomplete segment).

### Phase synchrony

To assess within-frequency phase synchrony, we used the weighted phase lag index (wPLI) (58), a measure of phase synchrony that de-weights zero phase (and antiphase) connectivity. For every 60 s segment and frequency band, raw wPLI between seed channel *j* (HPC) and target channel *k* (NC) was calculated as:

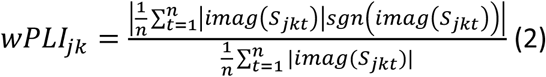

where *imag* indicates the imaginary part, *S*_*jkt*_ is the cross-spectral density between signals *j* and *k* at sample *t*, and *sgn* indicates the sign. We further created a normalized version of this metric using a surrogate approach. Surrogates were constructed by repeatedly (n = 100) time shifting the phase time series of the seed channel by a random amount between 1 and 59 s, and recalculating wPLI for each iteration. Note that time shifting is a more conservative approach than fully scrambling time series, which may result in spurious effects (78). This distribution was then used to z-score the raw wPLI value. Thus, the z-scored measure (wPLI_Z_) indicates how far, in terms of standard deviations, the observed coupling estimate is removed from the average coupling estimate under the null hypothesis of no coupling. We used the median to further aggregate wPLI and wPLI_Z_ values across data segments.

### Cross-frequency phase-amplitude coupling

For every 60 s segment, PAC was determined between all pairs of modulating frequency *f1* and modulated frequency *f2*, where *f2*>2\**f1*. We employed an adaptation of the mean vector length method (79) that adjusts for possible bias stemming from non-sinusoidal shapes of *f1* (60). Specifically, complex-valued debiased phase-amplitude coupling (dPAC) was calculated as:

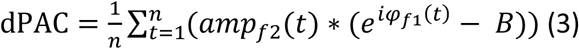

where *i* is the imaginary operator, *t* is time, *amp*_*f2*_*(t)* is the magnitude of the convolution result, or amplitude, of *f2*, *φ*_*f1*_*(t)* is the phase of *f1*, and *B* is the mean phase bias:

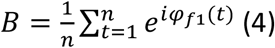

For same-site PAC (i.e., within HPC or within NC) *φ*_*f1*_ and *amp*_*f2*_ stemmed from the same electrode, whereas cross-site PAC used phase information from one brain structure and amplitude information from the other. Raw coupling strength (i.e., the degree to which the *f2* amplitude is non-uniformly distributed over *f1* phases) was defined as the magnitude (i.e., length) of the mean complex vector. For every 60 s segment, frequency pair, and same/cross-site combination we constructed a surrogate distribution of coupling strengths by repeatedly (n = 100) time shifting the *f1* phase time series with respect to the *f2* amplitude time series, and recalculating the mean vector length for each iteration. We then z-scored the observed coupling strength with respect to this null distribution of coupling strength values. We used the median to further aggregate dPAC_Z_ values across data segments.

### Cross-frequency modulation of phase synchrony

A modulation index (MI) was computed between all pairs of modulating frequency *f1* and modulated frequency *f2*, where *f2*>2\**f1*. For each frequency *f1*, samples were binned according to phase *φf1* (18 bins), and wPLI was calculated for each bin *b* and frequency *f2* following equation (2). Segment-averaged wPLI values were then used to calculate raw MI as:

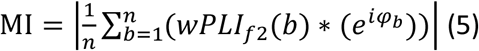

where *b* is the bin number, *n* is the number of bins (18), and *φ*_*b*_ is the phase at each bin center. This calculation was performed separately for HPC and NC *f1* phases. Note that MI was calculated across all available segments rather than per 60 s segment (as for wPLI and dPAC) because segment-wise MI estimates proved unstable. Similarly, surrogate distributions were constructed by repeatedly (n=100) shuffling the pairing of *f1* and *f2* phase segments (disallowing pairings where individual segments were unaltered), and recalculating MI across segments for each iteration, rather than per segment. Since the number of unique segment pairings depends on the number of available segments, 2 patients were excluded from N3 analyses. Observed MI values were z-scored with respect to their null distributions to generate normalized MI_Z_.

### Statistics

Statistical analyses were performed at both the group (wPLI/wPLI_Z_, dPAC_Z_, MI_Z_) and individual (wPLI_Z_, dPAC_Z_) levels. Phase synchrony (wPLI/wPLI_Z_) was assessed using non-parametric tests (Wilcoxon signed rank test or Mann-Whitney U test), followed by False Discovery Rate (q=0.15) correction for multiple comparisons (80). dPAC_Z_ and MI_Z_ were assessed using cluster-based permutation tests (81) with a *clusteralpha* value of 0.1. We used 1000 random permutations for most tests (i.e., N≥10), except for several cases where the number of possible permutations was lower (when N≤9), in which case each unique permutation was used exactly once. To determine the presence of effects, wPLI_Z_, dPAC_Z_, and MI_Z_ values at each frequency/frequency pair were compared to zero across patients (group) or data segments (individual) with one-tailed tests. Comparisons between sleep stages, regions, and directions were performed with two-tailed paired or unpaired tests as required. Clusters were deemed significant at P<0.05 (one-tailed) and P<0.025 (two-tailed). For dPAC_Z_ and MI_Z_, clusters were required to span at least 2 × 2 frequency bins.

### Data and code availability

Data are not publicly available due to privacy concerns related to clinical data, but data and accompanying analysis code are available from the corresponding or senior author upon obtaining ethical approval.

