## Supplementary Information for "Phase-based coordination of hippocampal and neocortical oscillations during human sleep"

#### Supplementary Results

##### Raw traces and spectrograms

Fig. S1 shows 10 s of concurrent scalp, HPC and NC traces and spectrograms for a single patient during N2, N3, and REM sleep. Scalp observations (top) were in line with typical sleep, with prominent N2 fast spindles, N3 SO/delta activity, and mixed frequency activity during REM. Unlike scalp spindles, HPC and NC spindles were poorly visible in the raw traces, but spectrogram activity showed distinct increases in spindle power. In contrast, HPC and NC often showed clear bouts of theta activity during N2 that were less obvious at the scalp. Also visible are occasional bursts of high-gamma activity in NC during N3. For virtually all frequency bands, activity across brain regions showed variable relations, with specific events (e.g., SO, theta, spindle) sometimes co-occurring, but mostly being expressed locally.

##### Power spectra within hippocampus and neocortex

HPC and NC power spectra were determined for each sleep stage, adjusting for 1/f power scaling with a slope-fitting procedure to enhance the visibility of narrowband peaks (Supp. Methods). Across individuals, adjusted HPC spectra (Fig. S2A) showed prominent spindle peaks and smaller delta/theta peaks during NREM. Small theta and spindle peaks were also visible during REM. All stages displayed a pronounced broad peak in the gamma range, consistent with human ripple activity. In contrast, NC showed clear evidence for SO activity during N3, but only weak NREM spindle peaks (Fig. S2B), consistent with Fig. S1 and previous reports of low spindle counts in temporal cortex (1). In addition, clear theta activity was present during N2 and REM, but not N3.

##### Individual phase synchrony profiles

Connectivity profiles for three example patients are shown in Fig. S3ABC. Single-subject analyses assessing above-chance phase coupling (one-tailed Wilcoxon signed rank test vs. zero) and stage differences (two-tailed Mann-Whitney U test) indicated quite heterogeneous effects. For example, p9 exhibited a multitude of peaks in the SO, delta, beta, gamma, and ripple (>85 Hz) ranges (Fig. S3C), phenomena not reliably observed across the full sample. In contrast, spindle synchrony peaks were present in all example patients for both N2 and N3, while clear theta N2/REM theta synchrony was present in two, further supporting the group analyses. Also note N3 beta synchrony at a faster frequency than where spindle power peaks for p9.

##### Relation between phase synchrony and power

Even with metrics like $wPLI_Z$, it remains a possibility that interregional phase synchrony is influenced by power in the two respective brain structures. However, while patients' phase synchrony profiles showed some visual resemblance to their HPC and NC power spectra in terms of theta and spindle peaks (compare Fig. S3 panels ABC to DEF, respectively), detrended HPC power and synchrony profiles were not significantly correlated for any sleep stage (within-subject Spearman correlation coefficients compared to zero with two-tailed Wilcoxon signed rank test: all $P_{uncorrected}>0.55$). Findings for NC were similar (all $P_{uncorrected}>0.19$), indicating that individual synchrony profiles do not merely reflect power. To further examine this notion across subjects, we trained k-nearest neighbor classifiers on individuals' power spectra and tested them on phase synchrony profiles, and vice versa. Across all 12 comparisons (3 stages x 2 regions x 2 classifier directions), resulting identification rates did not exceed 20% (binomial test: all $P_{corrected}>0.63$, $P_{uncorrected}=0.07$ at 20%). Focusing specifically on the theta, spindle, and beta frequencies that showed highest connectivity (7.4, 13.6, and 28.2 Hz), we observed no significant correlations between spindle connectivity and HPC or NC power across subjects for any sleep stage (all $P_{uncorrected}>0.09$). For theta, a significant relation between NC power and synchrony was found during N2 (Spearman's Rho: 0.89; $P_{corrected}=0.008$), but not during other sleep stages or for HPC power (all other $P_{uncorrected}>0.07$). For beta, a relation was found between NC power and synchrony during N2 ($P_{uncorrected}=0.03$), even though no above-chance connectivity was found for this frequency during this sleep stage. Moreover, this association did not survive multiple comparison correction, nor did other tests for the beta frequency (all $P_{corrected}>0.18$). Overall, these analyses indicate that phase synchronization is largely dissociated from local power. Further taking into account the removal of volume conduction by the employed wPLI metric, these findings suggest genuine interregional coordination of theta and spindle rhythms between human HPC and NC.

<u>Local cross-frequency coupling within hippocampus and neocortex</u>

*Group-level analyses*

Assessing the presence of PAC within brain structures (i.e., using phase and amplitude information from the same electrode), we observed systematic cross-frequency coupling for several frequency pairs, primarily during NREM. Within HPC, NREM PAC was most pronounced for SO-delta, SO-theta, SO-spindle, SO-ripple, and delta-ripple pairs (Fig. S4A). During REM, weak coupling was seen between the SO frequency and faster activity in the delta and theta ranges, consistent with the presence of limited SO activity in this sleep stage (2,3). These observations are qualitatively similar to our previous report using a larger number of HPC contacts (4).

A similar pattern of results was obtained for NC (Fig. S4B), again with a particularly strong organizing role of NREM SOs on several frequency bands, particularly delta and ripple activity. In contrast, spindles were only found to be weakly modulated by SOs during N3. In addition, we observed an orchestrating role of N2 theta on beta/gamma activity. During REM, we observed a weak but reliable modulation of ripple-band activity by frequencies ranging from the SO to theta bands. Direct comparisons between local HPC PAC and local NC PAC did not yield consistent differences (Fig. S5). In sum, for both HPC and NC separately, spectral components are robustly coupled, particularly during NREM.

*Patient-level analyses*

Besides assessing local PAC within HPC and NC at the group level, local PAC was also evaluated at the single-patient level (Fig. S6). Similar to group effects, PAC in both brain structures was apparent during NREM, but generally weaker during REM. As for the group effects, single-patient observations within HPC (Fig. S6A) closely mirror our previous results of individual variability (4), and are therefore only described as they relate to NC PAC.

For NC (Fig. S6B), we observed that the SO phase coordinates faster activity across a wide range of frequencies. Specifically, these include delta (p7, p9), theta (p2, p7, p9), spindle (p9), and beta/gamma/high-gamma components (p2, p7, p9), with modulations typically being stronger in N3 than N2. Interestingly, although SOs in both HPC and NC coordinated faster activity, the precise frequencies being modulated often varied between brain structures, even within individuals (e.g., p7, p9).

We also observed several instances in which neocortical activity was modulated by the NREM theta phase, with modulated ranges comprising the beta/gamma/high-gamma (p7, N2/N3), and beta (p9, N2) bands. Note that these effects often coincided with large theta peaks in the power spectrum. In contrast, the presence of a spectral theta peak was not invariably associated with fast activity being coupled to the theta phase (p2, N2). Moreover, spectral theta peaks could be present in both HPC and NC, while only showing theta coupling effects in NC (p7, N2), further emphasizing the differences in oscillatory organization between these brain areas. While theta-based modulation of faster activity was also seen during REM for one example patient (p7), these effects were not consistently present in the larger sample.

##### Interregional cross-frequency coupling between hippocampus and neocortex

###### *Patient-level analyses*

Similar to group-level cross-regional PAC as presented in Fig. 3AB, individual patients showed modulation of NC activity by HPC SOs during N3 (Fig. S8A), whereas the reverse was not the case (Fig. S8B). Other HPC-NC effects were heterogeneous, such as SO-spindle PAC during N3 (p9), and HPC theta modulation of NC beta-to-high-gamma activity during N2 (p7). NC-HPC PAC was also sometimes observed (e.g., N2 SO-theta and SO-ripple, and N2/N3 delta-ripple for p9).

#### Supplementary Methods

##### Spectral analysis

Procedures for spectral analysis were identical to our previous report (4). For each trial and channel, we estimated power spectral density using Welch's method with 3 s windows and 80% overlap (0.244 Hz resolution). Mean stage spectra were determined with a weighted average approach using trial durations as weights. Next, we removed the spectra's 1/f component to better emphasize narrowband spectral peaks. To this end, we first interpolated the notch-filtered region (50, 100, 150, and 200 Hz, ± 5 Hz) of each spectrum (Modified Akima cubic Hermite algorithm). Then, we fitted each N2 spectrum according to $af^b$ using log-log least squares regression (5,6). Fitting range was restricted to the 4–175 Hz range to avoid the often observed flattening of the spectrum below ~4 Hz and the ~200 Hz notch-interpolated data. Then, for each channel and stage, the N2 model fit was subtracted from the observed spectrum. We applied the N2 fit to all stages rather than using stage-specific fits to enable direct stage comparisons. Adjusted spectra were resampled to log space and smoothed three times with a moving average window of length 5.

#### Supplementary Figures

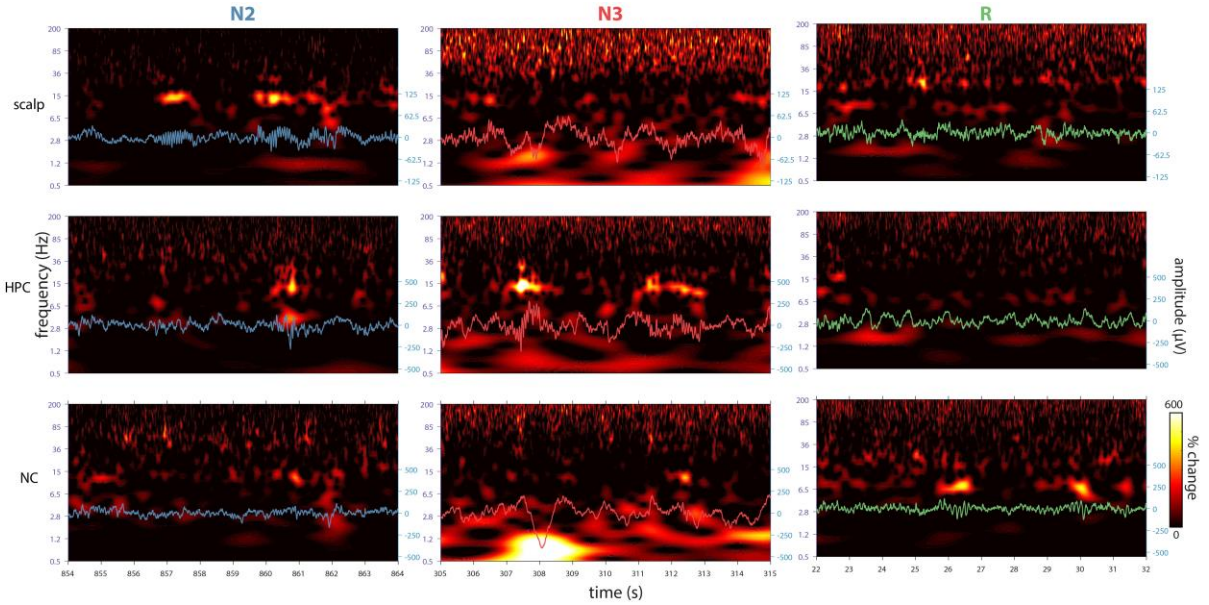


**Figure S1. Examples of concurrent sleep electrophysiology across brain structures.** Shown are 10 s segments of artifact-free data from a single patient (p2; see Fig. S3D for corresponding power spectra). Scalp, HPC, and NC traces show distinct features during each sleep stage, with scalp spindles not being matched by HPC or NC spindles (N2), nonsynchronous SO activity across sites (N3), and site-specific HPC delta and NC theta activity during REM. Note the different amplitudes of scalp and intracranial traces. Spectrograms show percent amplitude change relative to mean across all artifact-free N2, N3 and REM sleep.

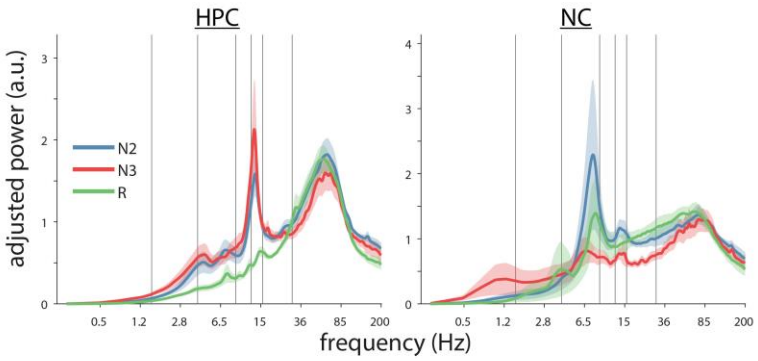


**Figure S2. Power spectra in hippocampus and neocortex.** Group-level spectra averaged across all 10 patients in HPC (A) and in NC (B). Error shading: standard error of the mean across patients. Gray vertical lines at 1.5, 4, 9, 12.5, 16 and 30 Hz indicate approximate boundaries between SO, delta, theta, slow spindle, fast spindle, beta, and faster activity.

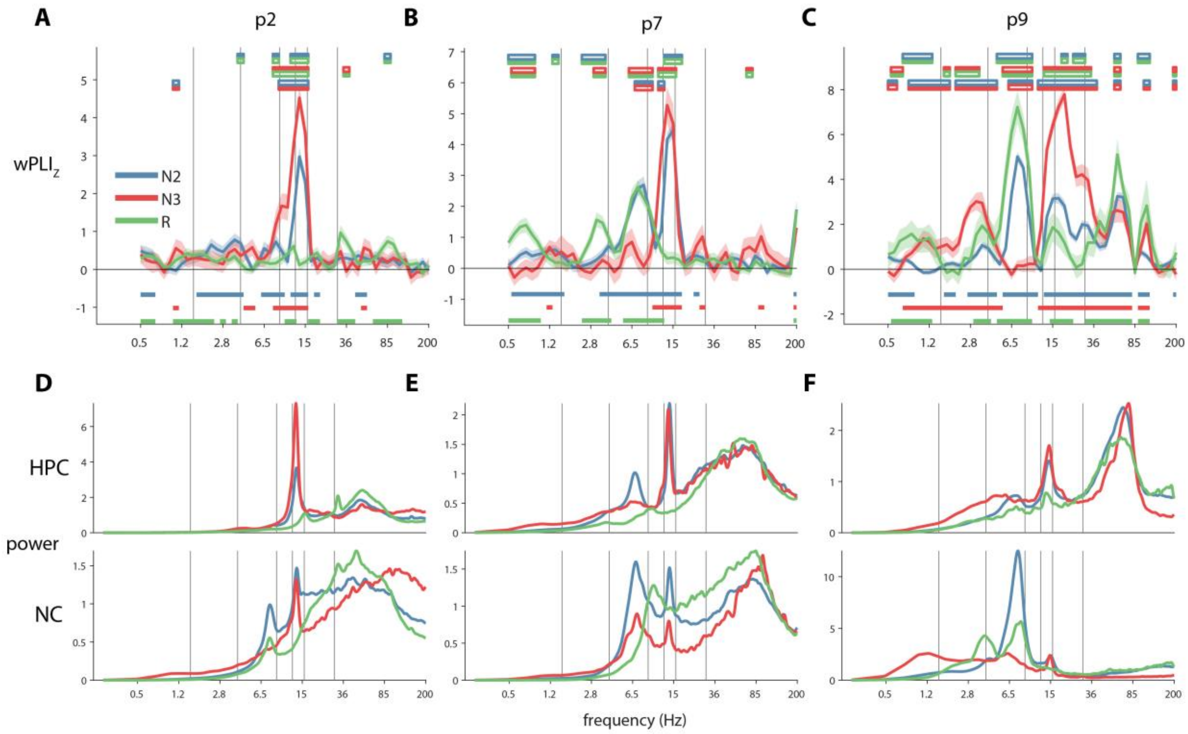


**Figure S3. Individual phase synchrony and power profiles.** (ABC) Example single-subject wPLI$_Z$ connectivity profiles, with stage comparisons (two-tailed Mann-Whitney U test with FDR) and comparisons to zero (one-tailed Wilcoxon signed rank test with FDR) indicated with color bars as in Fig. 2. (DEF) Corresponding power spectra in HPC and NC. Note the different scales for each panel.

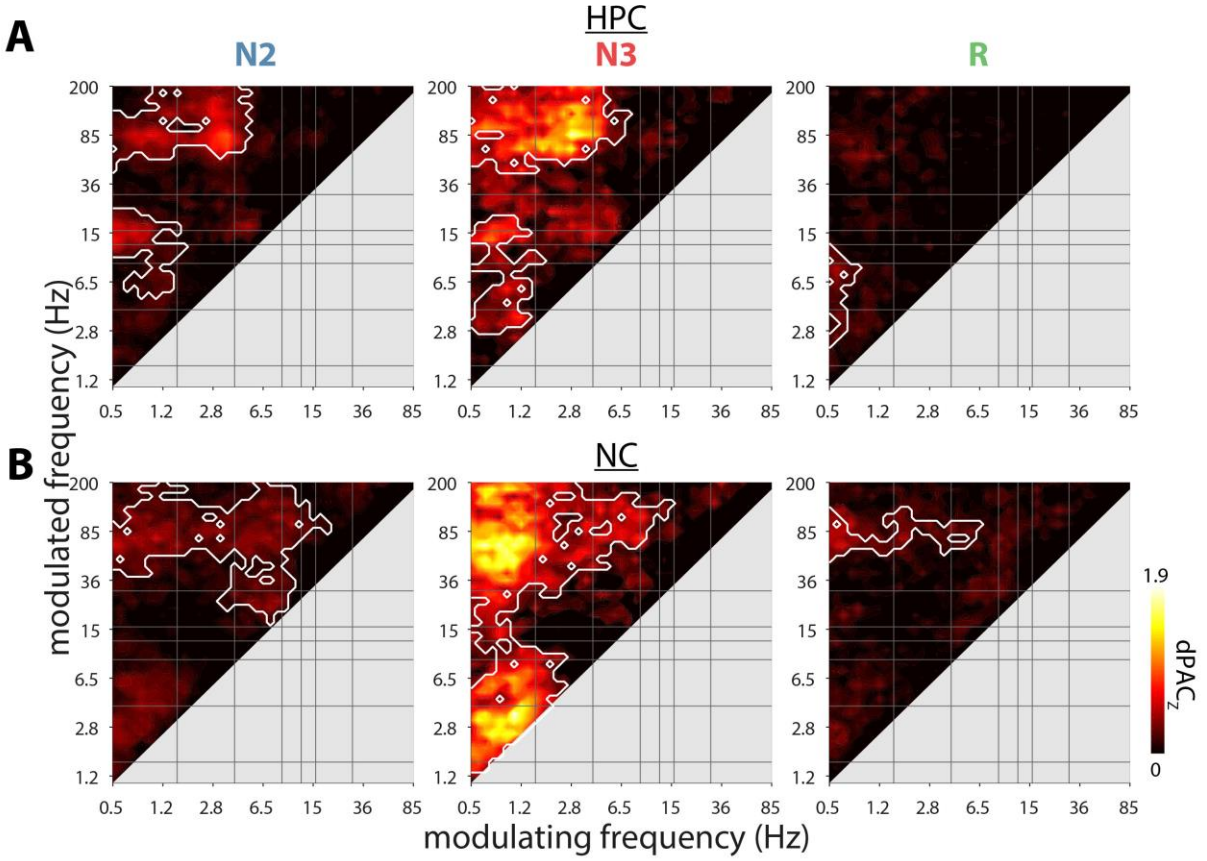


**Figure S4. Group-level cross-frequency coupling within hippocampus and neocortex.** Coupling strength profiles for HPC (A) and NC (B). White outlines indicate clusters of significantly higher than zero coupling across patients (cluster-based permutation tests).

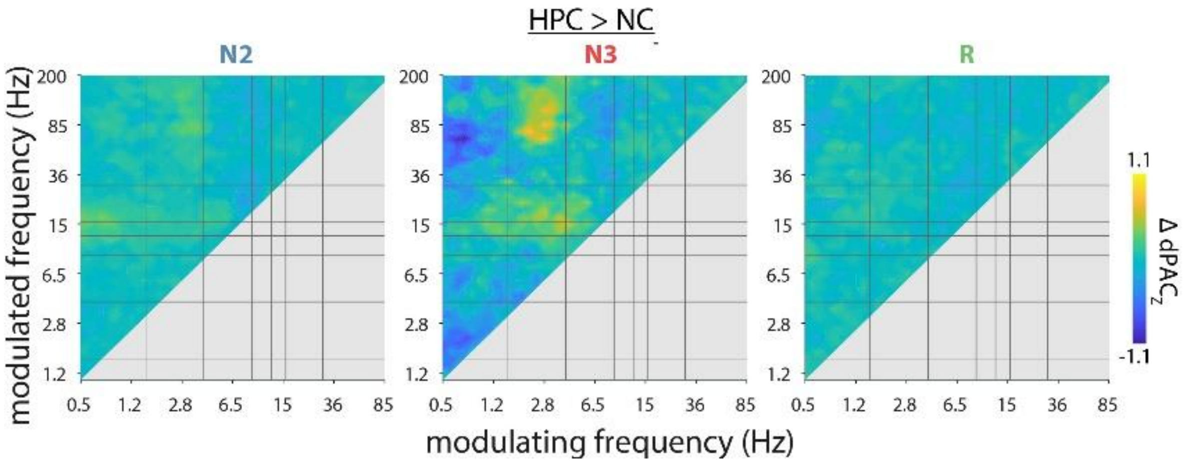


**Figure S5. Difference in local coupling strength between hippocampus and neocortex.** Although some differential coupling appears in N3, no clusters of systematically different PAC were found in either direction (two-tailed cluster-based permutation test).

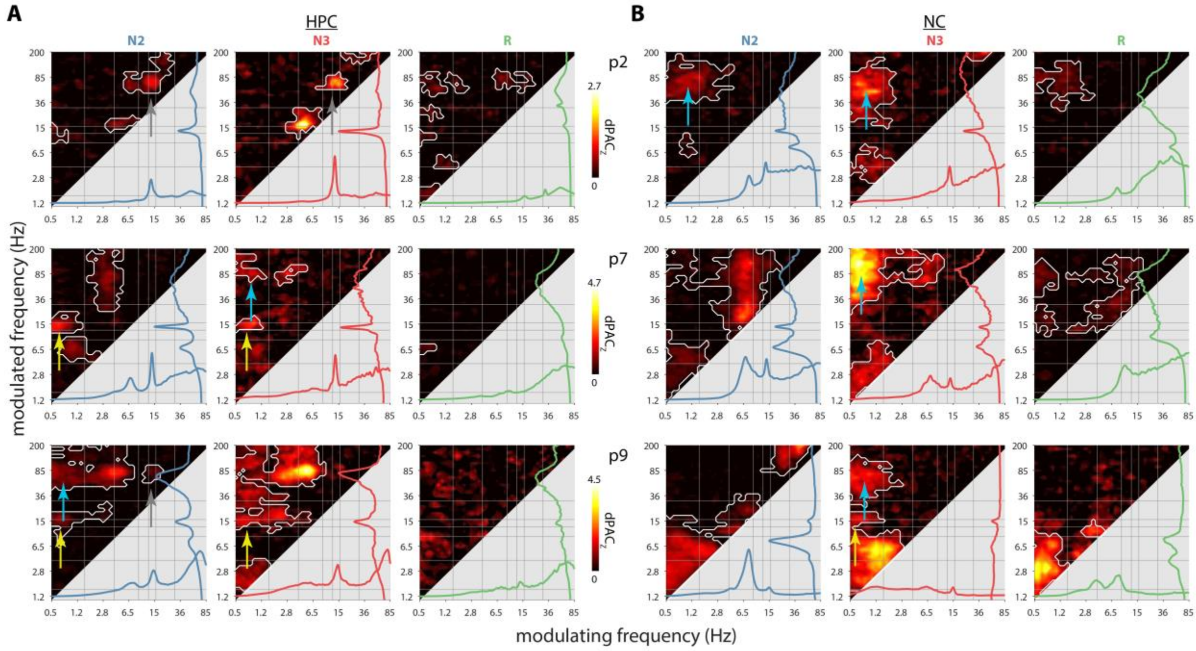


**Figure S6. Patient-level cross-frequency coupling within hippocampus and neocortex.** Coupling strength profiles for HPC (A) and NC (B). White outlines indicate clusters of significantly higher than zero coupling across one-minute data segments (one-tailed cluster-based permutation test vs. zero). Power spectra at the lower and right margins of each panel to provide a visual impression of the relation between PAC and power. Arrows indicate NREM SO-spindle (yellow), spindle-ripple (gray), and SO-ripple (blue) clusters.

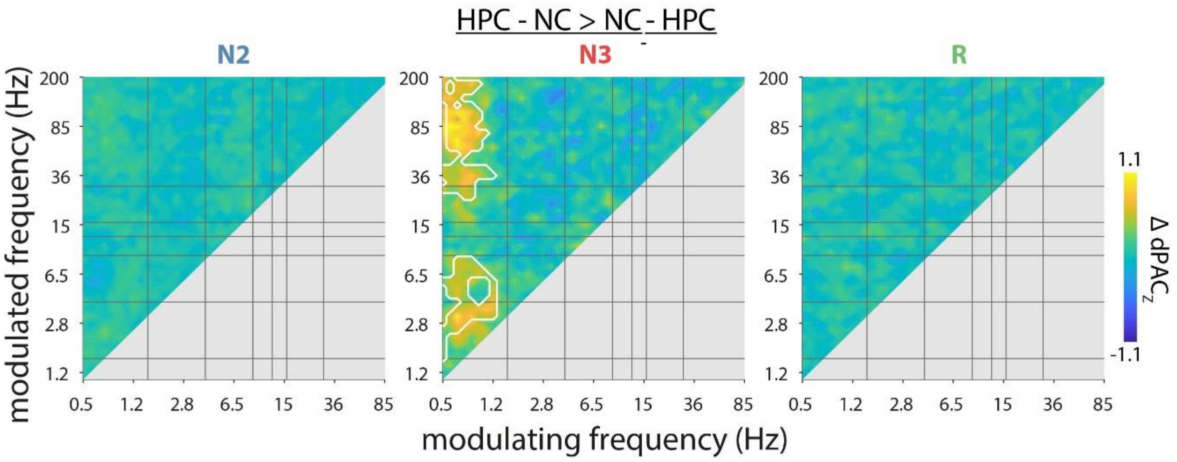


**Figure S7. Difference in interregional coupling strength between HPC-NC and NC-HPC directions.** White outlines indicate clusters of significantly greater local HPC-NC than NC-HPC PAC (two-tailed cluster-based permutation test). No clusters with greater NC-HPC than HPC-NC coupling were found.

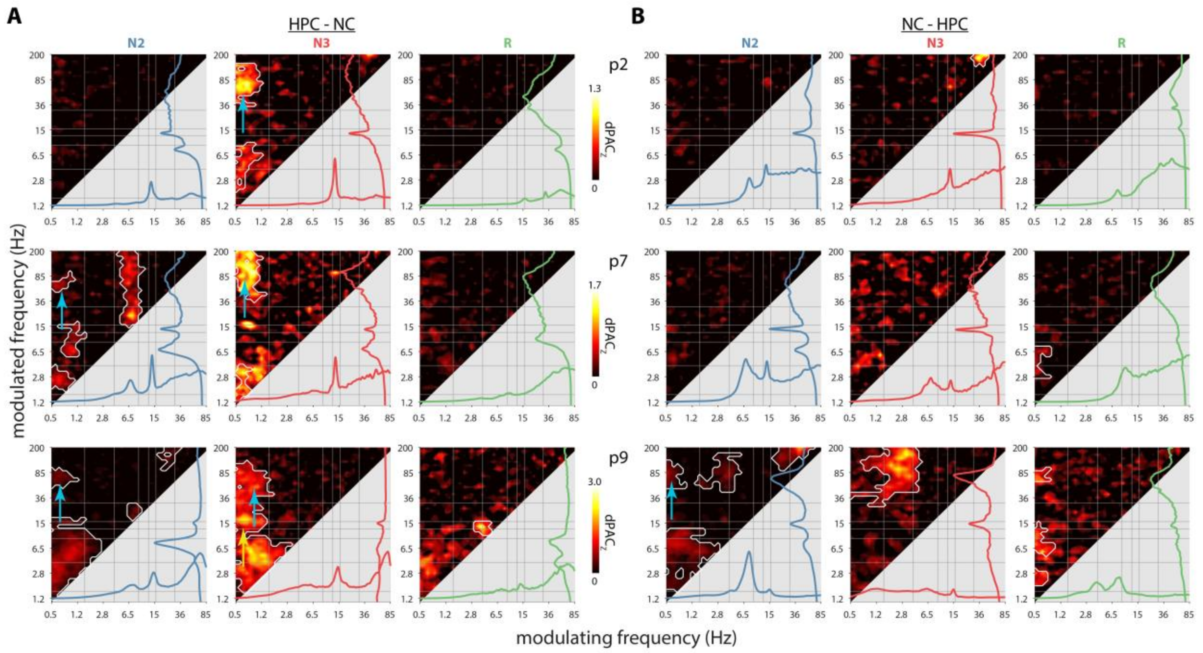


**Figure S8. Patient-level cross-frequency coupling between hippocampus and neocortex.** Coupling strength profiles for HPC-NC (A) and NC-HPC (B) for same patients as Fig. S6. White outlines indicate clusters of significantly higher than zero coupling across one-minute data segments (one-tailed cluster-based permutation test vs. zero). Spectra at the lower and right margins of each panel indicate power in the modulating and modulated region, respectively. Arrows indicate interregional NREM SO-spindle (yellow), spindle-ripple (white), and SO-ripple (blue) clusters.

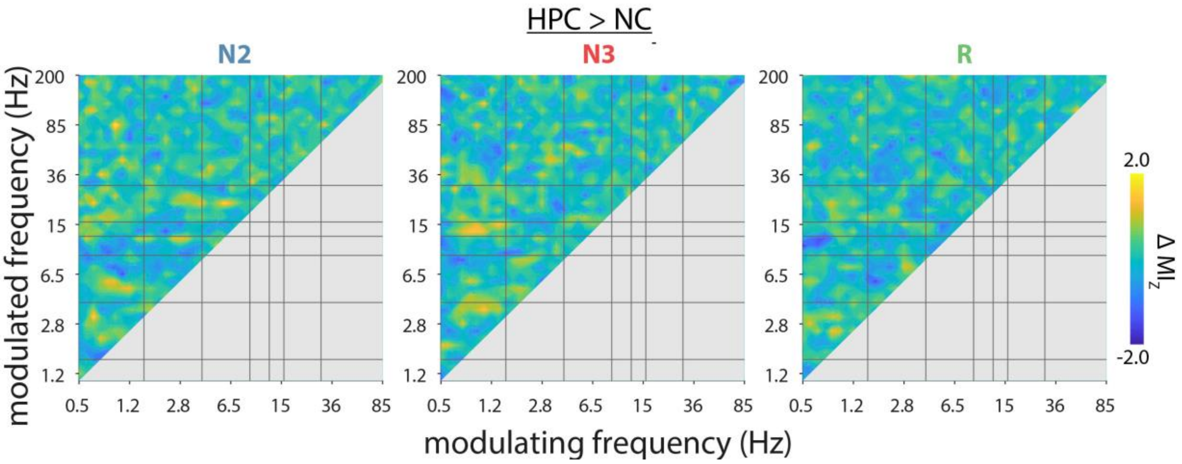


**Figure S9. Difference in cross-frequency modulation of phase synchronization between hippocampus and neocortex.** Cluster-based permutation tests revealed no clusters of differential modulation.

### Supplementary Tables

| | mean | SD |
|---|---|---|
| N1(%) | 32.3 | 23.1 |
| N2 (%) | 40.5 | 13.0 |
| N3 (%) | 12.3 | 7.6 |
| REM (%) | 14.9 | 7.0 |
| N1 (min) | 186.5 | 189.2 |
| N2 (min) | 198.1 | 66.7 |
| N3 (min) | 56.8 | 32.4 |
| REM (min) | 69.5 | 28.7 |
| total sleep (min) | 510.9 | 123.0 |
| WASO (min) | 96.8 | 77.5 |
| sleep efficiency (%) | 84.9 | 11.7 |

**Table S1. Sleep architecture.** WASO: wake after sleep onset.

| patient | sex | age (y) | epilepsy (y) | hemi | HPC | x | y | z | NC | x | y | z |
|---|---|---|---|---|---|---|---|---|---|---|---|---|
| p1 | M | 57 | 23 | R | P | 32 | -31 | -3 | MTG | 60 | -63 | 24 |
| p2 | M | 25 | 18 | L | M | -24 | -26 | -11 | STG | -71 | -34 | 8 |
| p3 | F | 62 | 49 | R | A | 29 | -11 | -20 | STG | 74 | -1 | -5 |
| p4 | M | 22 | 19 | L | P | -29 | -35 | -1 | STG | -62 | -28 | 11 |
| p5 | F | 44 | 29 | R | M | 27 | -33 | -4 | STG | 67 | -37 | 11 |
| p6 | F | 31 | 18 | L | P | -29 | -24 | -8 | MTG | -55 | -75 | 13 |
| p7 | F | 23 | 10 | R | P | 29 | -39 | -2 | MTG | 65 | -55 | 12 |
| p8 | M | 30 | 20 | R | P | 31 | -29 | -8 | MTG | 71 | -16 | -17 |
| p9 | M | 25 | 12 | L | A | -27 | -13 | -25 | MTG | -49 | -35 | -11 |
| p11 | M | 47 | 27 | L | A | -33 | -19 | -17 | ITG | -60 | -64 | -15 |

**Table S2. Patient and electrode details.** Selected HPC contacts were located in anterior (A), middle (M), or posterior (P) thirds of HPC, and NC contacts were located on inferior (ITG), middle (MTG), or superior (STG) temporal gyrus. xyz coordinates indicate electrode position after transforming to Montreal Neurological Institute template.

#### Supplementary Information References
